# Conscious perception of flickering stimuli in binocular rivalry and continuous flash suppression is not affected by tACS-induced SSR modulation

**DOI:** 10.1101/788950

**Authors:** Georg Schauer, Carolina Yuri Ogawa, Naotsugu Tsuchiya, Andreas Bartels

**Author notes:** Corresponding Author: Georg Schauer [ ], Address at which the study was carried out and present work address of corresponding author: Vision and Cognition Lab, Centre for Integrative Neuroscience, University of Tübingen Otfried-Müller-Straße 25, 72076 Tübingen, Germany.

## Abstract

The content of conscious perception is known to correlate with steady-state responses (SSRs), yet their causal relationship remains unclear. Can we manipulate conscious perception by directly interfering with SSRs through transcranial alternating current stimulation (tACS)? Here, we directly addressed this question in three experiments involving binocular rivalry and continuous flash suppression (CFS). Specifically, while participants (N=24) viewed either binocular rivalry or tried to detect stimuli masked by CFS, we applied sham or real tACS across parieto-occipital cortex at either the same or a different frequency and phase as an SSR eliciting flicker stimulus. We found that tACS did not differentially affect conscious perception in the forms of predominance, CFS detection accuracy, reaction time, or metacognitive sensitivity, confirmed by Bayesian statistics. We conclude that tACS application at frequencies of stimulus-induced SSRs does not have perceptual effects and that SSRs may be epiphenomenal to conscious perception.

## 1. Introduction

When participants are presented with a flickering visual stimulus, the firing rate of neurons in the early visual cortex synchronises to the frequency of the stimulus flicker and its harmonics (Regan, 1989; Herrmann, 2001; Pastor et al., 2003; Norcia et al., 2015, Vialatte et al., 2010). These oscillatory brain responses can also be recorded non-invasively by electroencephalography (EEG) and are known as steady-state responses (SSRs). SSRs are especially useful for tracing the perception of flickering stimuli by use of frequency tagging, where the presence of the stimulus’ frequency in the neural recording may indicate perceptual processing and even conscious perception of that stimulus (Andersen et al., 2011).

One issue that remains unclear is the causal role of SSRs. While they can correlate with conscious perception, and have been used to decode these (Brown & Norcia, 1997; Tononi, et al., 1998), it is unknown if they are causally involved in determining the content of conscious perception, or merely correlate with it. If they are epiphenomenal, they behave like a shadow does in relation to the object that cast it: while any change to the object leads to a change in the shadow, this relationship is purely unidirectional. Conversely, for neural oscillations not to be epiphenomenal to a process, then the process should be altered by interference with the oscillation (Sejnowski & Paulsen, 2006). Hence, if interfering with SSRs leads to a behavioural effect, then that activity can be said to be causally involved in bringing about that behaviour. The question is hence: can conscious perception be modulated by interfering with SSRs?

As Ruhnau et al. (2016) reported, SSRs can be modulated by the use of transcranial alternating current stimulation (tACS), a non-invasive brain stimulation technique. tACS interferes with neural oscillations through the application of a weak sinusoidal electric current on the head through two electrodes (Antal & Paulus, 2013; Herrmann et al., 2013).

tACS has been shown to affect conscious perception (Kanai et al., 2008; Laczó et al., 2012; Neuling et al., 2012a; Strüber et al., 2014) and behaviour (Pogosyan et al., 2009). Even though it is not completely understood how tACS interacts with neural oscillations, entrainment of neural oscillations is the prime candidate for explaining the mechanism behind the effect on neural oscillations during tACS stimulation (Neuling et al., 2012a; Witkowski et al., 2016; for opposing views see Battleday et al., 2014; Keitel et al., 2014). Entrainment here means the temporal phase alignment of neural oscillations with an external driving source (Thut et al., 2011; Vossen et al., 2015).

Ruhnau et al. (2016) argue that SSRs and tACS might be a natural fit, since the stationary sinusoidal structure of SSRs lends itself to the similarly stationary sinusoidal tACS modulation. They hypothesised that since alpha power increases with tACS, so should the SSR. Indeed, they showed that the amplitude of higher order harmonics of the SSRs can be increased when tACS at the same frequency as the SSRs is applied. They also demonstrated that a different frequency tACS had no discernible effect on the SSRs. They did not however aim to demonstrate that a modulation of the SSRs translates into a modulation of conscious perception. This is what we attempt to do in the present study. Two psychophysical paradigms lend themselves for testing this proposition.

Binocular rivalry occurs when two different images are presented to the two eyes in the same retinal location. Instead of seeing both images merged, participants report a spontaneous and unpredictable alternation between the two stimuli, irregularly spaced over time, where one image fades into awareness at the expense of the other (Blake & Logothetis, 2002). When SSR eliciting stimuli are used in binocular rivalry, the power of the SSRs has been demonstrated to correlate with conscious perception (Brown & Norcia, 1997). The amplitude of SSR is larger when the stimulus eliciting it is consciously perceived (Lawwill & Biersdorf, 1968). Tononi et al. (1998) used MEG frequency tagging and measured the SSRs for both of the stimuli’s frequencies. They found that frequency power was modulated contingent on perceptual dominance (Zhang et al., 2011).

Continuous flash suppression (CFS) is a variant of binocular rivalry, in which one eye is presented with the target stimulus and the other is presented with flashing dynamic high-contrast pattern masks (Tsuchiya & Koch, 2005). The dynamic Mondrian masks in one eye suppress perception of the target in the other eye, which remains invisible for dozens of seconds depending on the target contrast until it overcomes suppression and breaks into consciousness (Jiang et al., 2006; Yang et al., 2014). Crucially, CFS has been successfully employed at around 10 Hz (Tsuchiya & Koch, 2005), which is a frequency band also usable for recording SSRs.

Here we applied tACS to participants either while they viewed binocular rivalry or during a perceptual discrimination task under CFS in three separate experiments. The main objective of the present study was to find evidence for a causal role of SSRs on flickering perception by means of tACS. In other words, we intended to elucidate whether interference with SSRs can modulate conscious visual perception.

Specifically, two stimuli presented to each eye flickered at either 7.2 Hz (slow) or 9 Hz (fast). In binocular rivalry, each eye’s stimulus had one speed. For CFS, the mask was fast, while the target was slow. tACS parameters were applied in a three-by-two design: There were three tACS frequencies: slow, fast or sham. Sham tACS entailed no stimulation during experimental trials. However, to control for the sensation of tACS, which is strongest while tACS is ramped up and down prior and after each trial, we applied 8.1 Hz (midpoint between fast and slow) only during the ramp periods. Slow and fast tACS could also be either in-phase with the stimulus, where SSRs and tACS waveforms were in synchrony (0° phase lag), or out-of-phase, in which there was a 90° phase lag between the two waveforms. Since there is a lag between flickering stimulus presentation and the SSR onset, we performed an EEG pre-experiment which enabled us to estimate the SSR onset in order to consequently control the tACS phase to either 0 or 90 degrees. No EEG was recorded in the tACS experiment.

In the binocular rivalry experiment, we aimed to test if tACS frequency and tACS phase influenced percept dominance durations. Regarding the tACS frequency condition, we hypothesised that when tACS was set at the slow frequency, the stimulus flickering slowly would be perceived longer than the one flickering at the fast frequency, i.e. concurrent tACS should increase the SSR amplitude of the corresponding stimulus, making it more likely to enter consciousness. This is because the two SSR frequencies fluctuate according to the conscious perception of the participant. Increasing the SSR amplitude of the slow stimulus over the fast should lead to SSRs that are more characteristic of when the slow stimulus is consciously perceived. The same holds for the fast stimulus, while we would not expect an effect for the sham tACS condition. Regarding the tACS phase, we hypothesised that in-phase tACS (0° phase lag between SSR and tACS waveform) would increase predominance of the stimulus corresponding to the stimulation frequency, while the out-of-phase (90° phase lag) would not. We derived this hypothesis from previous findings of the effect of phase of ongoing EEG signals in particular frequency bands on behaviour: it has been demonstrated that stimuli presented in-phase to endogenous neural oscillations are perceived better compared to stimuli presented counter-phase (Sherman et al., 2016; Montemurro et al., 2008; Busch et al., 2009; VanRullen et al., 2011; Spaak et al., 2014). In fact, while in-phase stimulation enhanced performance in a letter discrimination task, out-of-phase stimulation had an opposing effect (Polania et al., 2012).

In a first CFS experiment we aimed to test if tACS frequency and phase influenced the ability to detect targets breaking through CFS. Participants were asked to indicate the position of a target as soon as they saw it, allowing us to compute accuracy and reaction time (RT). In a second CFS experiment we wanted to test the effect of tACS on metacognition. To this end, we presented participants with a CFS masked target for a period of time too short for the target to break into awareness in every trial. Participants were then asked to make a judgement on the position of the target paired with a confidence judgement. This allowed us to compute metacognitive sensitivity, i.e. how interospectively aware of the quality of their visual information processing participants were (Persaud et al., 2007; Rounis et al., 2010). While TMS over frontal cortex has been implied to impair metacognitive sensitivity (Rounis et al., 2010, for opposing evidence using theta burst stimulation see Bor et al., 2017), it was our aim to uncover if tACS impaired or even improved metacognitive sensitivity based on its parameters.

Regarding the tACS frequency, since the mask flickered fast while the target stimulus slowly, we hypothesised that the target would be detected faster when the tACS was set at the slow frequency, i.e. when tACS and target frequencies matched. Conversely, we expected that when tACS frequency matched the mask frequency (9 Hz), mask perception would be favoured and thus target detection would be hindered. Moreover, we proposed that in-phase tACS would have a favourable effect on task performance, while out-of-phase tACS would not.

## 2. Materials and method

### 2.1 Participants

24 healthy volunteers with normal or corrected to normal vision took part in the tACS experiments (mean age = 26.58 yrs ± 10.1 s.d., 12 female, 12 male, 2 left-handed). Out of this sample, 10 participants took part in the EEG pre-experiment (mean age = 26.60 yrs ± 2.99 s.d., 3 female, 1 left-handed). All participants were screened to meet health guidelines for neurostimulation and gave written consent. The study was approved by the local ethics committee.

### 2.2 Visual stimuli and apparatus

All stimuli were presented on a 27 inch monitor (width = 602 mm, ASUS, Taiwan) operating at 144 Hz, on a grey background (half of maximum illumination). There was no natural light contamination nor room lighting. Participants’ head position was fixed by a head and chin-rest. Binocular rivalry between two stimuli was created with a mirror stereoscope: The two stimuli were presented on the two sides of the screen separated by a board. The stereoscope then projected the images into the same retinal space of the participant. The mirrors were carefully adjusted for each participant to achieve fusion of the fixation cross and lines. The distance between monitor and participant through the stereoscope was 700 mm. All stimuli were created and controlled by a stimulus computer (Ubuntu 17.10) running Psychtoolbox 3 for Matlab R2014a (Mathworks, USA). Participants’ button press responses were collected with an adapted numeric keypad with eight buttons (two columns, four rows).

#### 2.2.1 Binocular rivalry stimuli

Two circular flickering checkerboards (both 3.5 degrees visual angle in diameter) were presented to the same retinal space in both eyes through the mirror stereoscope: one was black and green while the other was black and red (*figure* 1a). The checkerboard flickered at a preassigned frequency (see session 2.3), where the flickering was created through alternating presentation of the circular checkerboard and its inverted image. This method has been used successfully to elicit SSRs (Regan, 1966). Moreover, the checkerboards rotated clockwise (36 degrees/s) to reduce adaptation. Since EEG was not recorded in the tACS experiment, we ignored the possibility of the rotation inducing low frequency oscillations. Around each checkerboard there was a fusion aid to assist in keeping the two stimuli overlapping, which was a black and white checkerboard frame with a width of 7 degrees visual angle. The initial screen presented before the trial was comprised of the fusion aid in addition to a red fixation cross at its centre. The presentation eye of the checkerboards (i.e. which eye was presented with which checkerboard) was determined randomly for each trial.

#### 2.2.2 CFS stimuli

Two images were presented to the same retinal location in both eyes, a CFS mask and a grey background on which a target appeared (half of maximum illumination), both delimited by a square black and white fusion aid (12 degrees visual angle in width). The 200 CFS masks were composed of a set of colourful overlaid circles of various sizes. The target background included a single target (0.7 degrees visual angle in diameter), which was a low-contrast dark and light checkerboard, which could either be 1.5 degrees visual angle to the right or left side of a fixation cross (*figure* 2). The target flickered at a preassigned frequency (see session 2.3), where the flickering was created through alternating presentation of the circular checkerboard and its inverted image. The CFS masks changed at a frequency of 9 Hz. The initial screen presented before the trial start was comprised of a fusion aid and the red fixation cross. To begin each trial, participants pressed either the lower left or right button on the numeric keypad. The eye that was presented with the target and the side of the target with respect to the fixation cross (left or right) were randomised across trials.

### 2.3 Experimental design

Ten participants came to the laboratory on two separate days: once for an EEG-pre-experiment and once for three tACS experiments. The rest of 14 participants only participated in the tACS experiments. No EEG was recorded during tACS experiment.

#### 2.3.1 EEG pre-experiment

An important issue when applying tACS concurrent with SSRs is the difference in time needed for both to have a neural effect. The expected waveform for SSR and tACS evoked responses is sinusoidal but not concurrent in onset. SSRs occur after a given latency relative to stimulus presentation due to neural signal conduction delays from the eyes to the visual cortex, whereas the neural effect of tACS should be instantaneous. To ensure that the neural signatures of both overlap, tACS must be initiated at this latency. To do that, we attempted to estimate this latency in an EEG pre-experiment.

Participants were seated in a dark room, asked to put their chin onto the chin rest and instructed to fixate on the red fixation cross that was stereoscopically presented. The stereoscope was adjusted until a binocular match was achieved and participants reported only perceiving one single cross. Following a button press, participants viewed the binocular rivalry stimuli (section 2.2.1) for 300 trials of between 2.8 s and 3.2 s. In each trial, after an initial jitter period of 0.8-1.2 s from the start trial button press, both circular checkerboards appeared for 2 s, after which the initial screen followed. The next trial began when the participant pressed a button. Both binocular rivalry stimuli flickered in synchrony at the same frequency. In half the trials this frequency was 7.2 Hz, in the other 9 Hz. The task was to passively fixate at the fixation cross from pressing the trial start button until the stimulus disappeared. Participants were encouraged to take a longer break every hundred trials. The experiment lasted about 30 min. EEG was recorded continuously during this time (see section 2.4).

#### 2.3.2 tACS experiment 1: Binocular Rivalry

The first tACS experiment utilised binocular rivalry. Participants were instructed to fixate on the red fixation cross. After a button press, participants viewed the binocular rivalry stimuli (section 2.2.1) for 12 trials of 90 s each (*figure* 1b). During this time, the red checkerboard flickered at 7.2 Hz (slow frequency), while the green checkerboard flickered at 9 Hz (fast frequency). These frequencies were chosen as they have been previously used to elicit SSRs (Norcia et al., 2015), and also do not have matching harmonics below 35 Hz. Note that the precise choice of flicker frequency was constrained by the refresh rate of our monitor of 144 Hz (for a remedy see Andersen et al., 2015). Participants were instructed to report their perception by pressing and holding one of two buttons using their right hand: the left button while the red checkerboard was dominant, and the right button for green. Moreover, participants were asked to press no button during perceptual mixtures.

tACS was used continually throughout the 90 s trials. tACS intensity was linearly ramped up (0 to 1000 mA) during 10 s towards the beginning of the trial, and ramped down after it ended. There were six experimental conditions in a three-by-two design: three tACS frequency conditions: 1) 7.2 Hz for the slow condition, 2) 9 Hz for the fast condition and 3) sham; and two phase conditions: 1) In-phase, where tACS onset was the SSR onset minus the ramp-up time. 2) Out-of-phase, where tACS onset was the SSR plus 1/2 the period of the tACS frequency minus the ramp-up time. In the sham condition there was no electrical stimulation during the trial, although the current fade-in and out were present at 8.1 Hz (midpoint between the slow and fast frequencies) so that participants were not able to know that the simulation was off, since most of sensation of having tACS applied occurs during the fade-in and fade-out periods. After the fade-in, the tACS was shut off for the duration of the trial and then reengaged at the end at the same intensity to which the fade-in ramped up. Each of the three tACS frequency conditions (fast, slow, sham) appeared for four consecutive trials. Each of these four trials was randomly assigned to one of two tACS phase conditions (in-phase or out-of-phase with the stimulus). The order of conditions was randomised.

#### 2.3.3 tACS experiment 2: CFS - RT

The second tACS experiment utilised CFS. Participants were instructed to fixate on the red fixation cross. Following a button press, participants viewed the CFS stimuli (section 2.2.2) for 12 blocks of 90 s each. Each block had 16 trials. In each trial, the stimuli were presented for 5 s, during which the circular checkerboard (target) gradually faded in from 0 contrast to the maximal contrast over 1 second and the CFS mask gradually faded out from maximal contrast to 0 contrast over 1 to 5 second (*figure* 2a, inset). Since the target could be either to the left or right side of the fixation cross, participants were instructed to press the left or right button as soon as they saw the target. With this, we measured stimulus detection through accuracy and RT. The target flickering frequency was 7.2 Hz (slow frequency) while the CFS mask change rate was 9 Hz (fast frequency). After 5 s, identical CFS masks were presented to both eyes for 250 ms to reduce afterimages. Then the plain gray fixation screen was presented for 375 ms before the following trial began. Experimental conditions and tACS were identical to experiment 1 (*figure 1*; except eye 1 receives the slow target and eye 2 the fast mask).

#### 2.3.4 tACS experiment 3: CFS - Metacognition

The third tACS experiment was identical to the second one using the CFS stimuli (section 2.2.2), except for the following differences: In each trial, the stimuli were presented for 1 s only and the contrast of target and CFS mask remained constant. Following presentation of the mask for 250 ms to reduce afterimages (*figure 2b*). The target flickering frequency was 7.2 Hz while the mask change frame rate was 9 Hz. Afterwards, a grey screen with red fixation cross was displayed for 4 s. Participants were instructed to report the side of the target with confidence rating. To that end, participants had four right and four left buttons, to indicate the side as well as how sure they were on a 1-4 scale by pressing one of the eight buttons. Then the grey fixation screen was presented for 375 ms before the following trial began. Experimental conditions and tACS were identical to experiments 1 and 2 (*figure 1*; except eye 1 receives the slow target and eye 2 the fast mask).

### 2.4 EEG recording and SSR latency analysis

EEG data were recorded only during the EEG pre-experiment at 2500 Hz acquisition rate using a BrainAmp MR Plus system (Brain Vision; bandpass 0.016–250 Hz) with 11 active actiCap Ag/AgCl electrodes placed over occipital and parietal areas (in the international 10-20 system: Pz, P1, P2, POz, PO3, PO4, PO7, PO8, Oz, O1, O2) as well as the ground (AFz) and reference (FCz) electrodes, at an impedance lower than 5 kΩ (*figure* 1d). EEG data were processed offline with ERPLAB, a plugin for the EEGLAB toolbox in Matlab R2014a (Mathworks, USA). The signal was downsampled to 256 Hz, referenced to the average of all electrodes and bandpass-filtered with the cutoff values of 0.05 Hz and 30 Hz. Baseline-corrected epochs were extracted for time intervals of - 500 ms to +1000 ms around the 300 trial onsets in pre-experiment EEG session (figure 3a) as well as ∼4000 stimulus updates (target flicker) across all pre-experiment trials (figure 3b). This period was sufficient for detecting the SSR. The epochs were averaged for each participant and later into a grand average.

**Figure 1:**
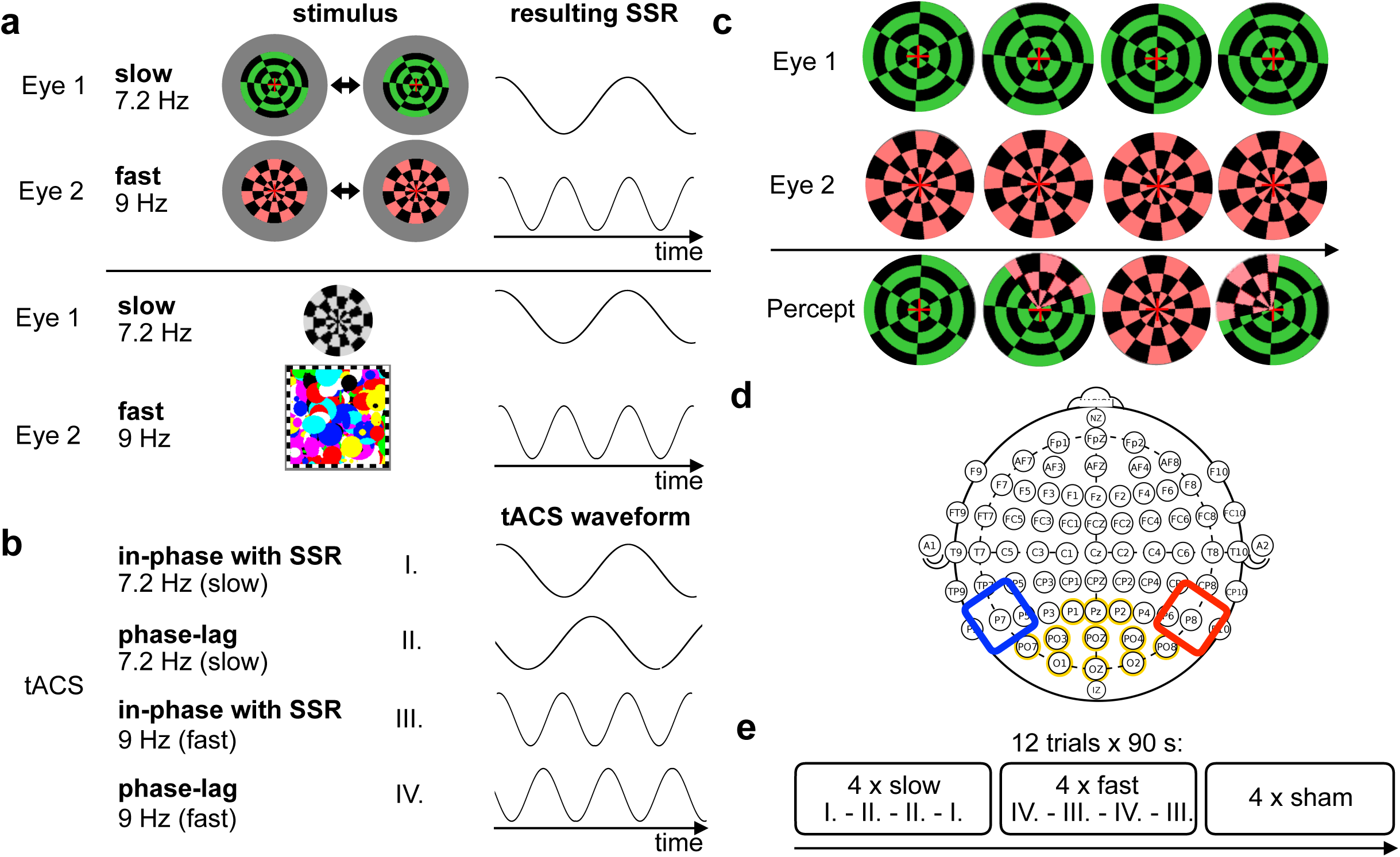
materials & tACS experiment 1. **(a)** Top: Each eye was presented with a different circular checkerboard (green and red, respectively) in the same retinal location, with a red fixation cross in the centre. Flickering emerged through alternating presentation of the circular checkerboard and its inverted image (see left and right) as a square wave. One eye receives a fast (9 Hz) the other a slow (7.2 Hz) flicker, leading to a corresponding SSR. Bottom: The target (here shown in Eye 1) consisted of a dark and light flickering circular checkerboard that could be to the left or right of the fixation cross (shown here in large as illustration). The CFS mask (here shown in Eye 2) was a series of images of colourful circles (200 different patterns). All stimuli were delimited by a square black and white fusion aid. The target flickered at 7.2 Hz alternating between its original and inverted pattern. CFS masks were updated at 9 Hz. **(b)** tACS could be slow (I. & II.) or fast (III. & IV.) also. Within each, tACS can be applied in-phase (I. & III.) or out-of-phase with lag (II. & IV.) **(c)** Illustration of binocular rivalry stimuli and percept evolution over time. Both eyes were presented with either of the two checkerboards (upper two rows), which led to a percept that fluctuated between the two eyes (bottom row). Stimuli rotated clockwise at 36 degrees/s. Participants reported percept types by pressing and holding one of two buttons throughout 90 s of viewing time per trial. **(d)** Montage overview: Yellow are electrodes that were used for recording EEG in the pre-experiment within the international 10-20 system. Blue/red are cathode and anode for the tACS placed over the P7/P8 electrode sites. **(e)** Design overview: There were 12 trials of 90 s. tACS fast, slow and sham appeared four times each in consecutive blocks, counterbalanced. Within these, tACS was applied twice in-phase, twice out-of-phase (or 90 deg phase lag), counterbalanced.

**Figure 2:**
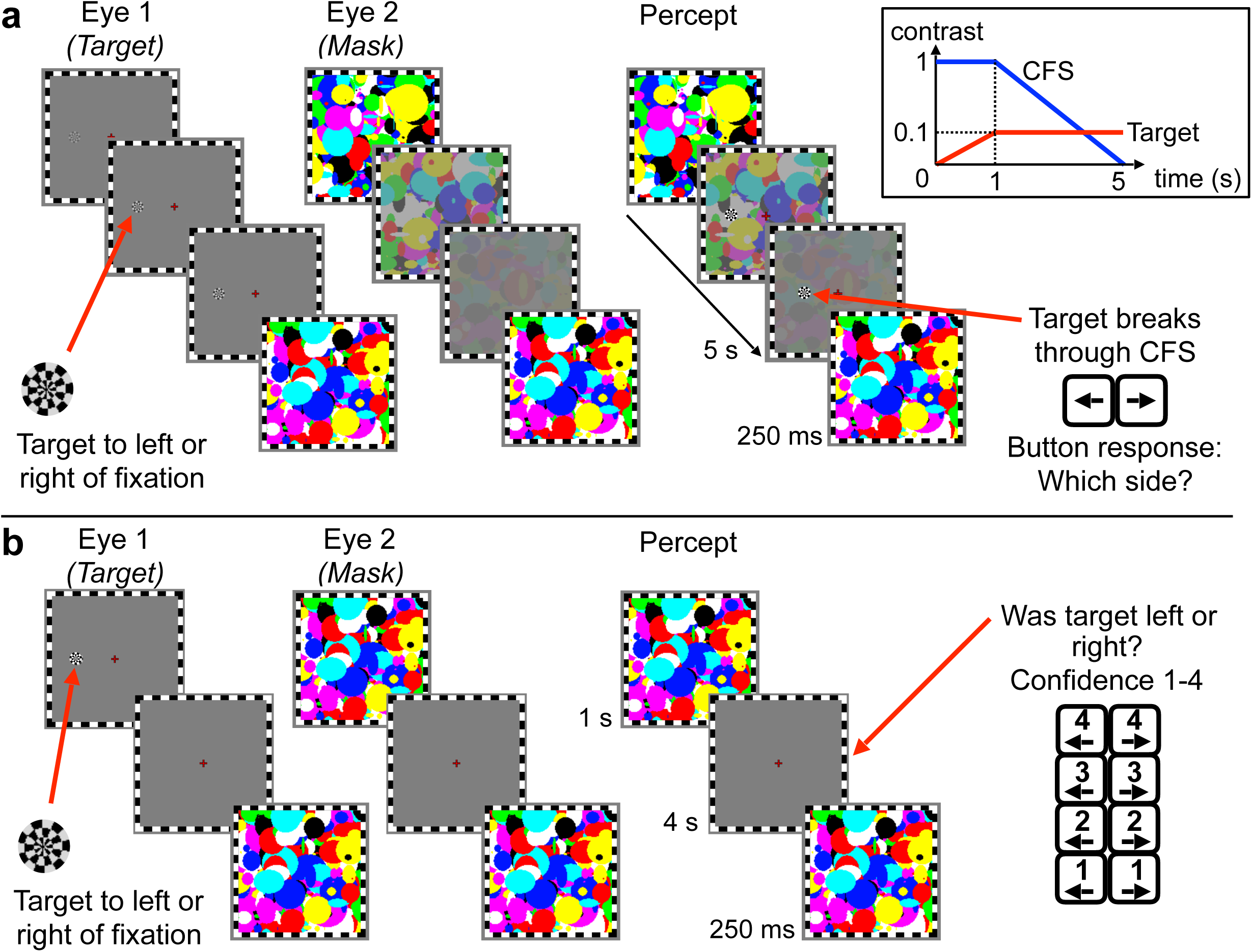
tACS experiments 2 & 3. The target canvas (here shown in Eye 1) consisted of a black and white flickering circular checkerboard that could be to the left or right of the fixation cross (left shown here). For illustration, the target (shown on the left here) is displayed at a contrast level of 10% of maximum. The CFS mask (here shown in Eye 2) was a series of images of colourful circles (200 different patterns). All stimuli were delimited by a square black and white fusion aid. **(a)** Experiment 2: In each trial, the stimuli were presented for 5 s during which the CFS mask gradually faded out (at maximum contrast for 1 s and then reducing to 0 linearly for 4 s) and the checkerboard gradually faded in (from 0 to 10% maximum contrast in the first second). At some point within these 5 s, participants were likely to see the target break through the mask. Following the task, the mask was presented to both eyes for 250 ms to reduce afterimages of the target. **(b)** Experiment 3: In each trial, the stimuli were presented for 1 s. Then participants had 4 s to make a judgment paired with a confidence rating. Following the task, the mask was presented to both eyes for 250 ms to reduce afterimages of the target.

**Figure 3:**
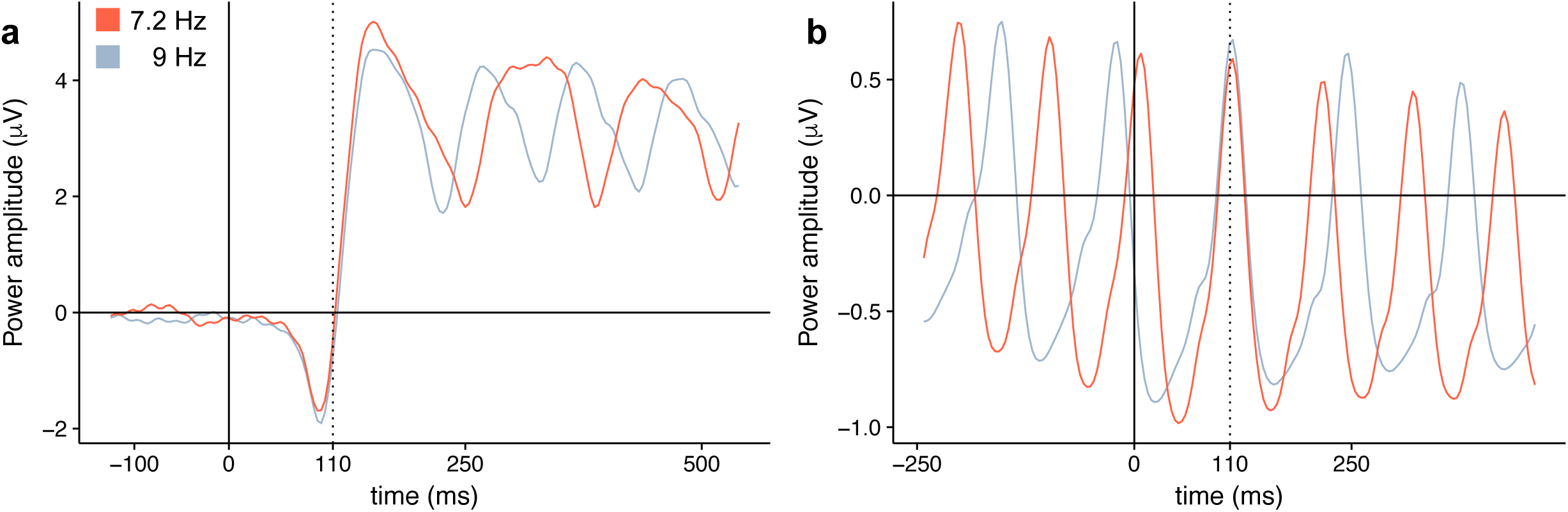
EEG signal traces of the pre-experiment. **(a)** Baseline-corrected grand averaged onsets of SSRs for 10 participants, showing traces evoked by the fast stimulus (9 Hz flickering frequency) in red and by the slow stimulus (7.2 Hz flickering frequency) in blue for the Pz electrode. Stimulus onsets were at time 0. The EEG signal fell into the repeating SSR pattern beyond 110 ms. **(b)** Grand averaged EEG signal over the O2 electrode, where time = 0 represents the onset of each stimulus flicker (or refresh). At about time = 110 ms, the two signals peaked.

**Figure 4:**
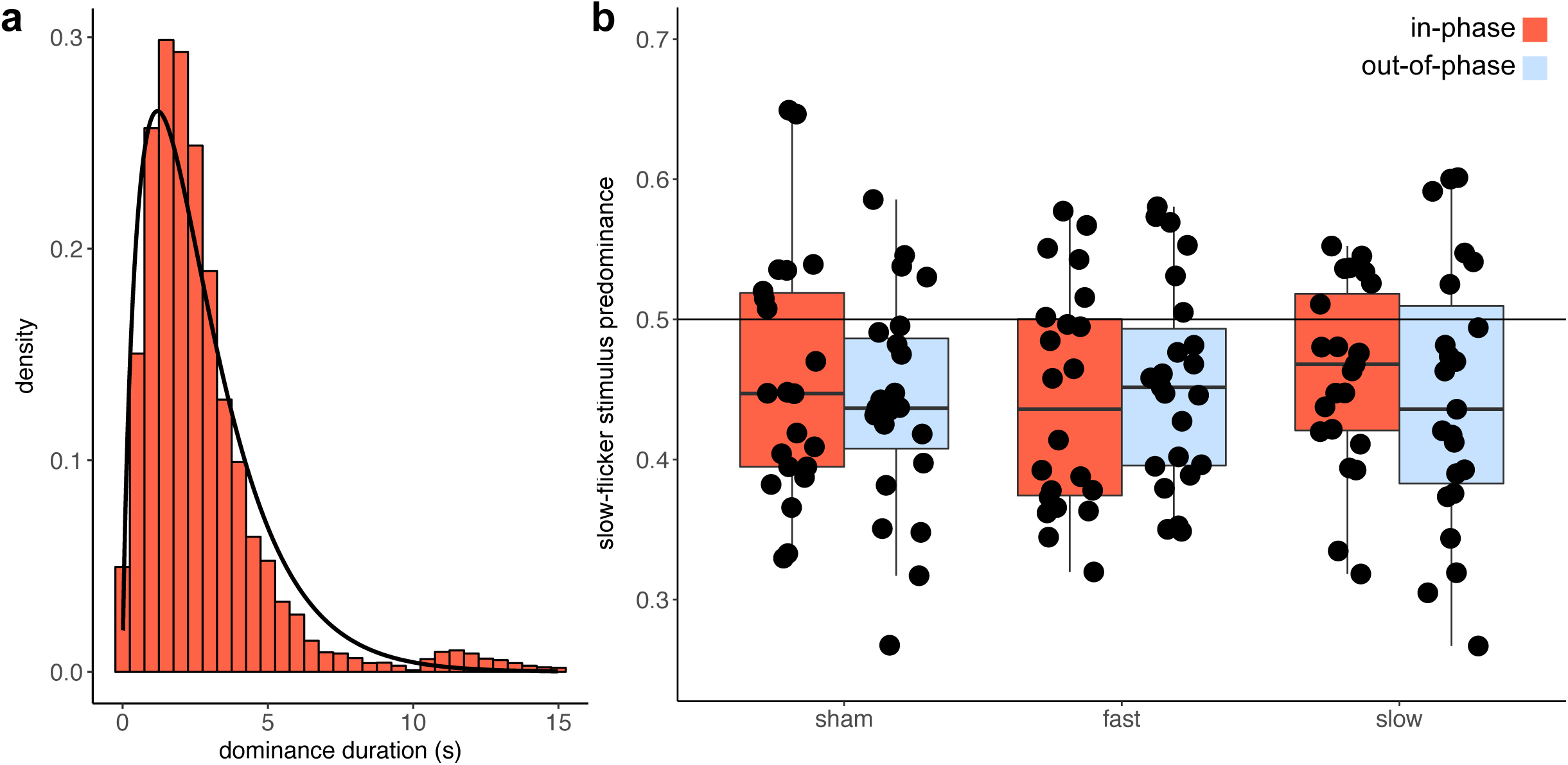
tACS experiment 1 results. **(a)** Binocular rivalry dominance duration histogram (pooled from all trials of all participants), with density distribution overlaid. The durations follow a gamma distribution (shape = 1.78, scale = 0.65, median dominance is 2.61 s). **(b)** Predominance across conditions (N = 23): The slow predominance (y axis) represents the proportion of the sum of all slow (red target) dominance compared to the sum of all perceptual dominances (slow and fast). If predominance is 0.5, then both percepts were perceived in equal sduration. Each dot is a data point of one participant, separately for tACS frequency condition (sham, fast, slow) and tACS phase condition (in-phase: red, out-of-phase: blue). Boxplot of median encased by the 25th and 75th percentile.

SSRs offer good signal to noise ratios, as a large portion of signal power is contained within a few frequency bands. Since these are known to the experimenter, their isolation from the EEG signal is comparably easy, even by eye. Visual inspection of the SSRs elicited in the average of the 11 electrodes as well as separately electrode Pz for each participant individually, revealed that the delay between the stimulus being drawn on the screen and the specific SSR waveform changed its polarity at around 110 ms after the onset of trial (*figure* 3a). In addition, our flicker reversal-related ERP shows the peak positivity at around 110 ms after the onset of both the slow and fast flicker (figure 3a) (Please remember that during this pre-experiment binocular rivalry, the flicker rate to both eyes was the same (see section 2.3.1)). This pattern was observed across all participants. This is important, because in order to phase-lock the tACS to the SSRs, the tACS onset must be adjusted by this SSR onset.

### 2.5 tACS parameters

The stimulus computer controlled the battery-driven tACS device (DC-Stimulation Plus, NeuroConn GmbH, Ilmenau, Germany). tACS is mainly driven by the parameters of frequency, current amplitude/intensity and stimulation phase (Antal & Paulus, 2013). An alternating sinusoidal current at 1000 µA intensity was delivered through two conductive-rubber electrodes that were positioned on the scalp at P7 and P8 positions according to the International 10-20 system (*figure* 1d). Prior studies have shown that perceptual and neural effects depend on tACS intensity: 500 μA or higher of 10 Hz tACS over visual cortex (electrodes 4 cm over inion and on vertex, corresponding to Cz/Oz) were sufficient to cause visual phosphenes in darkness (Kanai et al., 2008). 140 Hz tACS stimulation of M1 had an inhibitory effect on TMS induced MEP amplitude at 400 µA, had no effect at 200 µA, 600 µA and 800 µA, but had an excitatory effect at 1000 µA (Moliadze et al., 2012). We therefore chose 1000 µA to form part of an excitatory protocol.

The electrodes were placed into sponges (5.5 x 6 cm size) and soaked in 0.9% NaCl solution, resulting in impedances lower than 20 kΩ. In all experiments, tACS was applied for 90 s at a time (see session 2.3). Right before the beginning and right after the end of each trial, the current linearly faded-in (from zero to 1000 µA) and out (from 1000 µA to zero), respectively, during 10 sec.

### 2.6 Behavioural data analysis

#### 2.6.1 Experiment 1 (binocular rivalry)

Our measure of interest was the modulation of predominance as a function of the different tACS conditions. To that end, we first extracted rivalry dominance durations from the button presses of participants. We excluded button presses that were interrupted by the end of the trial and treated double button presses and no button presses as mixed percepts. We next investigated if the data followed previously reported gamma distributions (Levelt, 1967) for each participant and each condition separately. Predominance was calculated as the sum of all slow (red target) dominance durations divided by the sum of both (green and red targets) dominance durations. We used this measure for our main analysis, since it indicates if an experimental manipulation biased participants to see the slow over the fast stimulus contingent on how tACS was applied. Finally we entered predominance as dependent variable into a repeated measures ANOVA, using tACS frequency and phase as factors.

Next to predominance, we decided to also test whether tACS had modulated dominance durations overall, as well as for only the fast stimulus, slow stimulus and mixes percepts. We therefore entered each of these as dependent variable into separate repeated measures ANOVAs, using tACS frequency and phase as factors.

#### 2.6.2 Experiment 2 (CFS RT)

We were interested in the effect of tACS on RT and accuracy. For each participant and each condition separately, we first examined if accuracy was higher than 0.8 to ensure participants adequately performed in each condition. Then we looked at all RTs to see if they followed a gamma distribution. The median RT for all trials within a condition was then used for the main analysis. Here we entered RT and accuracy as dependent variables into two separate repeated measures ANOVAs, using tACS frequency and phase as factors.

#### 2.6.3 Experiment 3 (CFS meta-cognition)

As a measure of detection and metacognitive accuracy, we used the area under curve (AUC) of the receiver operating characteristic (ROC) in type I and type II, respectively. To obtain type I ROC curves, the data was grouped into 8 levels according to judgement and confidence with 7 partitions. Starting with a threshold where -4 indicates *p1*, and all other buttons *p2*, we calculated a hit and false alarm rate. Next this can be done assuming that -4 and -3 indicate *p1* and all others *p2*, and so forth. Hence, for each criterion, we obtain a set of numbers, that can be plotted against each other, yielding the ROC curve. The area under this curve now represents discrimination performance, where 0.5 is chance and larger areas mean better detection sensitivity (Sherman, et al. 2015). To obtain type II ROC curves, the data was grouped into 4 levels according to correctness of judgement and confidence. First, let 1 represent low confidence and 2-4 high, then 1-2 represent low and 3-4 high, and so forth. For each criterion, we calculated the type II false alarm rate, which is the proportion of high confidence trials, when the participant is incorrect, as well as the hit rate, which is the proportion of high confidence trials, when the participant is correct. Connecting these points yields the type II ROC curve. To this end, we used the method and code provided by Fleming & Lau (2014). The advantage of this non-parametric method is its robustness against violations of Gaussian equal variance. The area under this curve now represents metacognitive sensitivity (Matthews et al., 2018; Matthews et al., 2019).

Finally, we entered type I AUC and type II AUC as dependent variables into two separate repeated measures ANOVAs, using tACS frequency and phase as factors.

## 3. Results

### 3.1 Experiment 1: Binocular Rivalry

The dominance durations pooled from all participants together followed a gamma distribution (*figure* 4a). We therefore chose the median as a measure of central tendency. One participant, whose data was corrupted by faulty data recording, needed to be excluded from the analysis. The average of all participants’ median dominance durations was 2.58 s ± 0.83 s.d. for the fast flickering stimulus (green target), 2.24 s ± 0.74 s.d. for the slow stimulus (red target) and 2.39 s ± 0.74 s.d. considering both together.

Entering the predominances into a repeated measures ANOVA, using as factors tACS frequency (sham, slow, fast) and tACS phase (in-phase, out-of-phase), we found that there was neither a significant main effect of tACS frequency (F(2,44) < 1, *ns*), tACS phase (F(1,22) = 1.03, *ns*), nor a significant interaction between the two factors (F(2,44) < 1, *ns*) (*figure* 4b). Since the absence of an effect is not evidence in favour of the null hypothesis (H0), we used Bayesian statistics to ascertain how strong the above null results were. Using the BayesFactor package (0.9.2) for R64, we modelled a Bayesian ANOVA with the mathematical underpinnings presented by Rouder et al. (2012), computing a Bayes Factor (BF) for each combination of factors and interaction, against the null that all effects are 0. The analysis resulted in a BF of 0.09 (strong evidence for H0) for the main effect of tACS frequency, a BF of 0.2 (substantial evidence for H0) for main effect of tACS phase, and a BF of 0.002 (very strong evidence for H0) for the interaction term. This provides overall strong evidence that the tACS conditions did not affect predominance.

ANOVAs using other dependent variables, such as median dominance duration for fast, slow, mixed and all percepts, yielded similar null results.

### 3.2 Experiment 2: CFS - RT

The RT values pooled across all conditions and participants followed a gamma distribution (*figure* 5c). We therefore chose the median as measure of central tendency. The same participant as in experiment 1 needed to be excluded from the analysis due to data corruption. The average of all participants’ median RT was 2.01 s ± 0.47 s.d. Moreover, participants correctly identified the position of the target most of the time at an overall accuracy of 96%. For the subsequent RT analyses, incorrect trials were excluded.

**Figure 5:**
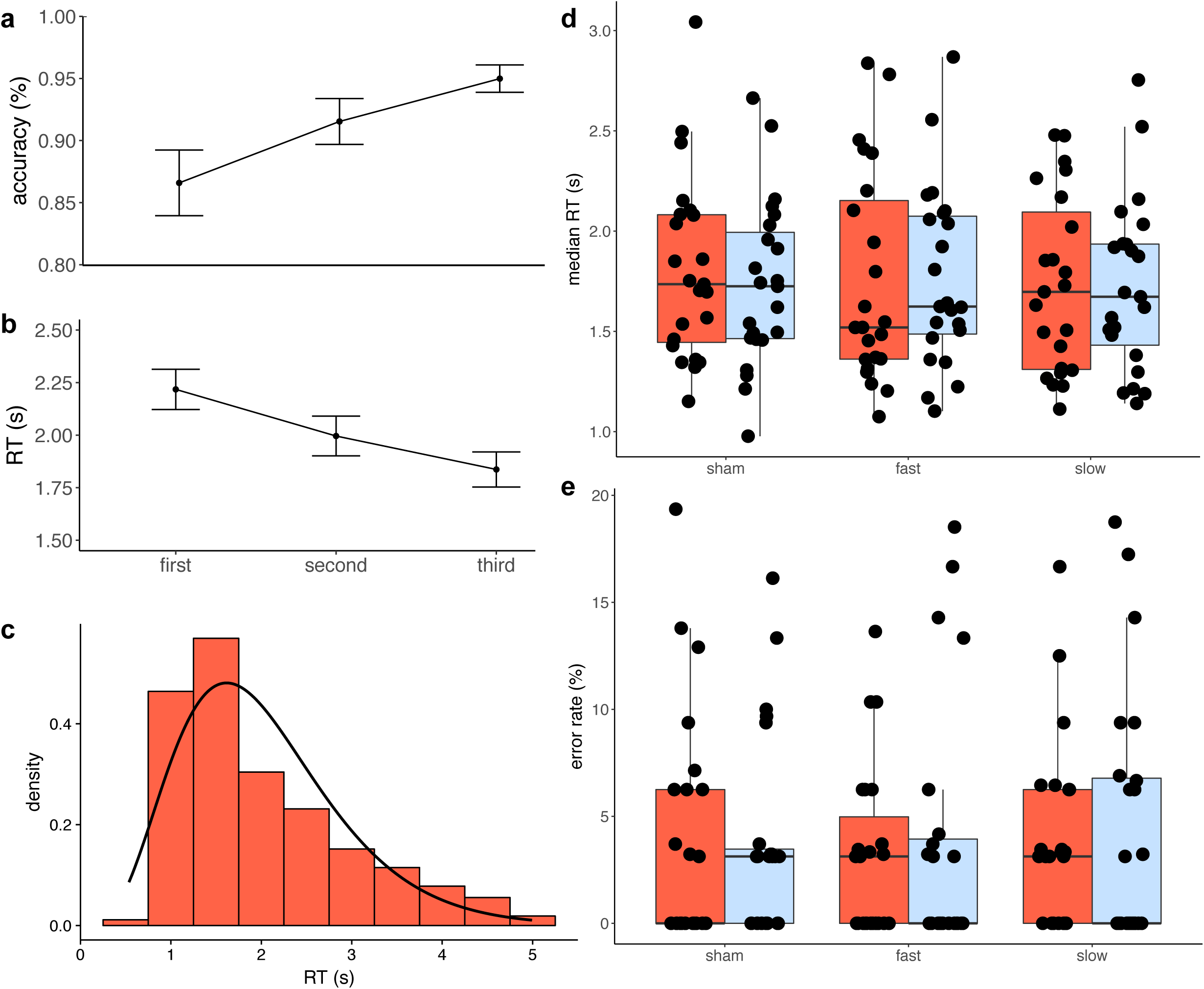
tACS experiment 2 results. Effect of time (block number) on accuracy **(a)** and median RT **(b):** On the x axis are the first, second and third experimental blocks according to temporal order and independent on what the tACS frequency condition was. Error bars indicate SEM. **(c)** RT histogram pooled from all trials of all participants (N = 8282). The durations follow a gamma distribution (shape = 4.96, scale = 2.45, median RT is 1.70 s). **(d)** Median RT and **(e)** error rate across experimental conditions (N = 23). Error rate was calculated as 100% - % correct. Each dot is a data point one participant, separately for tACS frequency and phase condition (In-phase: red, out-of-phase: blue). Boxplots of median encased by the 25th and 75th percentile.

**Figure 6:**
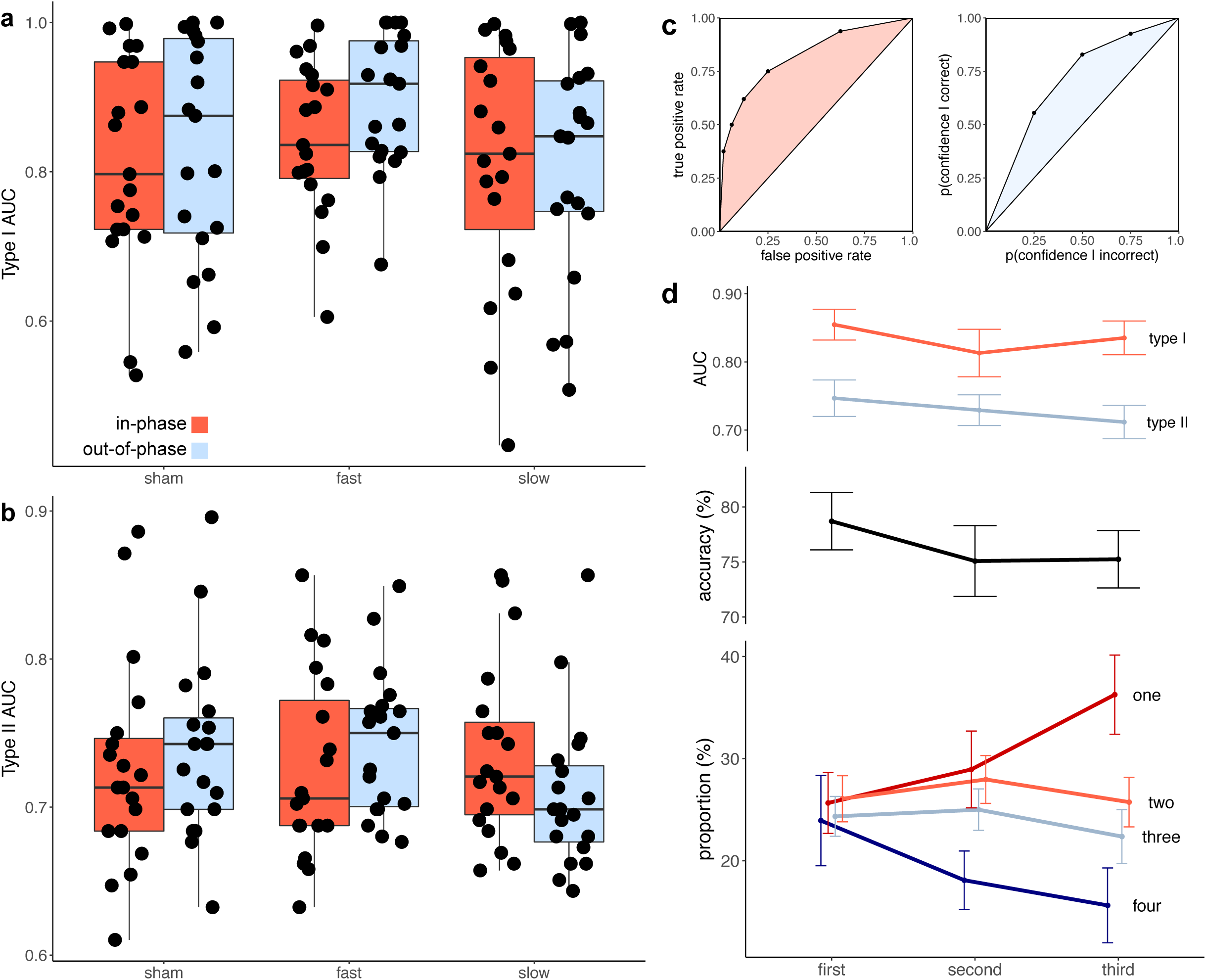
tACS experiment 3 results. **(a)** Type I AUC and **(b)** type II AUC across experimental conditions (N = 19). Each dot is a data point from one participant, separately for tACS frequency and phase condition (In-phase: red, out-of-phase: blue). **(c)** Left: Type I ROC for a representative participant. For each confidence criterion, type I false alarm rate (x) is plotted against type I hit rate (y). AUC. Right: Type II ROC curve for a representative participant. For each confidence criterion, type II false alarm rate (x) is plotted against type II hit rate (y). AUC, representing metacognitive sensitivity. Boxplots of median encased by the 25th and 75th percentile. **(d)** Effect of time (block number) on AUC (top), accuracy (middle), and the proportion of button presses corresponding to the different confidence levels from 1-4 (bottom). On the x axis are the first, second and third experimental blocks according to temporal order and independent on what the tACS frequency condition was. Error bars indicate SEM.

The median RTs for each condition from each participant were entered into a repeated measures ANOVA with two factors: tACS frequency (sham, fast, slow) and tACS phase (in-phase, out-of-phase). Mauchly’s test indicated that the assumption of sphericity had been violated for the interaction term (χ^2^(2) = 8.84, p = 0.01), therefore degrees of freedom were corrected using Greenhouse-Geisser estimates of sphericity (ε = 0.74). The ANOVA showed that there was neither significant main effect of tACS frequency (F(2,44) < 1, *ns*) nor tACS phase (F(1,22) < 1, *ns*), nor a significant interaction between the two (F(2,44) < 1, *ns*) (*figure* 5d). A Bayesian ANOVA produced a BF of 0.08 (strong evidence for H0) for the main effect of tACS frequency, a BF of 0.2 (substantial evidence for H0) for the main effect of tACS phase, and a BF of 0.002 (very strong evidence for H0) for the interaction term. This provides overall strong evidence that the tACS conditions did not affect RT.

When accuracy was entered into the same ANOVA as dependent variable, it showed that there was neither a significant main effect of tACS frequency (F(2,44) < 1, *ns*), tACS phase (F(1,22) < 1, *ns*), nor an interaction between the two (F(2,44) < 1, *ns*) (*figure* 5e). This null effect was again supported by Bayesian statistics, which produced a BF of 0.09 (strong evidence for H0) for the main effect of tACS frequency, a BF of 0.2 (substantial evidence for H0) for the main effect of tACS phase, as well as a BF of 0.002 (very strong evidence for H0) for the interaction term. This provides overall strong evidence that the tACS conditions also did not affect accuracy.

Despite the absence of a tACS effect, performance in the task could have improved over time due to perceptual learning or training. To test this, the three experimental blocks (comprised of four 90 s trials each) were sorted chronologically, regardless of the tACS frequency condition. To begin, we calculated a linear regression to test if the tested block was a significant predictor of accuracy and found a positive relationship, such that experience led to higher performance (Formula: Accuracy = 0.83 + 0.04 * block, F(1, 70) = 9.22, p = 0.003, R^2^ = 0.10; *figure* 5a). Next, we asked if time was also a significant predictor or RT and found a negative relationship, such that training led to a lower RT (Formula: RT = 2.40 – 0.19 * block, F(1, 70) = 8.80, p = 0.004, R^2^ = 0.10; *figure* 5b). Both these results together point out that more time elapsing led to better performance.

### 3.3 Experiment 3: CFS - Metacognition

Type I and II AUC were computed for each participant individually. A representative set of curves is shown in *figure* 6c. 5 participants needed to be excluded from the analysis for being at ceiling or floor in terms of confidence. The mean type I AUC for all participants was 0.83 ± 0.14 s.d., the mean type II AUC was 0.73 ± 0.06 s.d.

First, type I AUC for each condition from each participant was entered into a repeated measures ANOVA with factors tACS frequency (sham, fast, slow) and tACS phase (in-phase, out-of-phase). We found that the main effect of tACS frequency approached significance (F(2,36) = 2.87, p = 0.07), whereas there was no main effect of tACS phase (F(1,18) = 1.83, *ns*) nor interaction (F(2,36) < 1, *ns*) (*figure* 6a). A Bayesian ANOVA showed that there was only anecdotal evidence in favour of the main effect of tACS frequency (BF = 0.44). Visual inspection of the data revealed that this was merely due to less spread of the data in the fast condition. The analysis furthermore produced a BF of 0.28 with regard to the main effect of tACS phase (substantial evidence for the H0) and a BF of 0.022 for the interaction (very strong evidence for the H0). In essence, the analysis showed that tACS phase and its interaction with frequency had no modulating effect on type I AUC, while we cannot draw any conclusion about the main effect of tACS frequency.

Next, we constructed an equivalent ANOVA with type II AUC as dependent variable. We found no main effect of tACS frequency (F(2,36) < 1, *ns*), nor of tACS phase (F(1,18) < 1, *ns*). However, there was a significant interaction term (F(2,36) = 6.39, p = 0.005). This interaction indicates that tACS frequency modulated the effect of tACS phase, even in the absence of main effects. This can be visually seen in the data (*figure* 6b), where out-of-phase tACS produced a higher mean type II AUC in the fast and sham frequencies, while this effect was reversed for the slow frequency. However, already visual inspection cautions us for overinterpretation, since it could easily be attributed to noise, given the spread of the data. Bayesian statistics confirm this suspicion, yielding a BF of 0.01 for the interaction, which is substantial evidence for the null. Similarly, for the main effects of tACS frequency and phase, BF of 0.12 and 0.2 respectively also point towards the H0.

As in experiment 2, despite the absence of a tACS effect, performance in the task could have improved over time due to perceptual learning. To test this, the three experimental blocks (comprised of four 90 s trials each) were sorted chronologically, regardless of the tACS frequency condition. First, linear regressions were calculated to test if time was a significant predictor of the proportion of the different confidence levels. We found that the lowest confidence level (button 1) was pressed more often as time went on (Formula: Proportion = 0.20 + 0.05 * block, F(1,55) = 4.50, p = 0.04, R^2^ = 0.06). There was furthermore a trend for fewer presses of the highest confidence button as time progressed (Formula: Proportion = 0.27 - 0.04 * block, F(1,55) = 2.56, p = 0.12, R^2^ = 0.12). There was no trend evident for the middle two confidence levels.

Hence, the lowest confidence button was pressed 5% more each block, while the highest confidence button was pressed 4% less (*figure* 6d bottom). Despite training, this interestingly indicates less confidence over time. This is especially remarkable since accuracy did not change significantly (F(1,55) > 1, *ns*) (*figure* 6d middle). These results are captured also in the respective ROC curves: While there was no change in type I AUC (F(1,55) > 1, *ns*), metacognitive sensitivity decreased over time (Formula: AUC = 0.53 - 0.04 * time, F(1,55) = 4.13, p < 0.05, R^2^ = 0.05) (*figure* 6d top).

## 4. Discussion

In this study, we tested whether application of tACS at congruent or incongruent frequencies (or phases) with stimulus-induced SSRs influenced conscious perception. In particular, we used two types of stimuli that we expected to be highly sensitive for detection of neural intervention effects on conscious perception - binocular rivalry and continuous flash suppression - as both allow for continuous shifts of perceptual balance towards one or the other percept while keeping the physical input to the visual system equivalent between various tACS conditions. Effects on conscious perception, in the form of binocular rivalry dominances as well as target detection under CFS, would entail that modulation of the SSR causally influences perception (Sejnowski & Paulsen, 2006). That would be relevant since it could elucidate whether SSRs only correlate with visual perception, or if their modulation influence it. Our results, however, showed that the different tACS conditions did not affect the perceptual measures: predominance (in the binocular rivalry experiment), RT and accuracy (in the first CFS experiment) as well as detection and metacognitive sensitivity (in the second CFS experiment) were not significantly different across tACS conditions. tACS did not differentially affect binocular rivalry dominance durations, improve or impair detection of targets masked by CFS, nor modify metacognitive sensitivity. Thus, one might tentatively conclude that modulation of SSRs does not alter the dynamics of perception of flickering stimuli, but instead only correlate with them. What we observed were time-on-task effects independent of tACS. In the first CFS experiment, RT and accuracy improved over time, as could be expected (Zizlsperger et al., 2016). In the second CFS experiment, contrary to our expectation, metacognitive sensitivity decreased with time, an effect driven by a decrease in confidence. Several explanations could account for this: participants could have become more tired and less concentrated, making it harder to detect the near threshold stimuli, or, the lack of perceptual feedback led to increasing insecurity for psychological reasons. Alternatively, it may be that tACS had a cumulative negative effect on cognitive performance, however, this is unlikely given the positive time-on-task effect observed in the first CFS experiment. Instead, our participant instructions did not emphasise the equal distribution of confidence for a given block, meaning that what we measured may not actually have reflect trial-by-trial fluctuation of confidence, but instead meta awareness of fatigue, attentional drift or mindwandering over blocks (Matthews et al., 2018).

### 4.1 tACS modulation of SSRs

Concluding from our results that SSRs do not have any causal role in determining the dynamics of perception of flickering stimuli may be premature though: since we did not record EEG during the two main experiments, we cannot claim with certainty whether SSRs were effectively modulated by tACS in our study. Especially in the CFS experiments, where targets were not presented foveally, we cannot even say with certainty that the targets evoked an SSR. Moreover, Ruhnau et al. (2016a) reported SSRs modulation only at the 3f and 4f harmonics rather than at fundamental frequency and only when tACS was applied at the same frequency as the SSR. Conversely, tACS reduced phase synchrony at fundamental frequency and 2nd harmonic. If SSRs at the fundamental frequency are indeed not influenced by tACS, but only at its 3f or 4f harmonics, we should have chosen lower tACS frequencies in order to affect the fundamental SSR frequency.

Specifically, 3 Hz (9/3, as 3f harmonic) or 2.25 Hz (9/4, as 4f harmonic) for the fast condition, and 2.4 Hz (7.2/3, for 3f harmonic) or 1.8 Hz (7.2/4, for 4f harmonic) for the slow condition. This lack of tACS effect on the fundamental SSR frequency may however be due to the phase lag between tACS and SSR, which Ruhnau et al. (2016a) did not control for. Since in our experiment, the phase relationship between tACS and SSR was controlled, we believe it more likely that fundamental SSR frequencies were modulated by tACS. However, we must concede that without EEG evidence, we cannot be certain, especially since it is possible that the phase lag did not remain constant during stimulation. Still, even if we were only modulating higher SSR harmonics, (9 × 3 = 27 Hz and 9 × 4 = 36 Hz SSRs elicited by fast flickering stimulation; 7.2 × 3 = 21.6 Hz and 7.2 × 4 = 28.8 Hz elicited by slow), we would still have modulated the SSR (albeit only in part), and a positive result would have allowed us to make inferences as to our hypotheses. In this case, we may cautiously conclude that our results cannot demonstrate the absence of any causal role of the SSR, but rather the absence of a causal role of higher order dynamics. However, we are cautious about this conclusion, given that the mechanism leading to the result by Ruhnau et al. (2016a) remains elusive and is not supported by modelling work (Reato et al., 2010; Neuling et al., 2012b).

### 4.2 Potential issues of tACS orientation and location in eliciting SSRs

There are several other explanations for our null result though. To begin, our electrode montage was chosen based on the work of Neuling, et al. (2012b). By use of finite-element models to predict current flow through the brain during electrical neurostimulation dependent on electrode position, Neuling, et al. (2012b) were able to show that the P7/P8 montage was particularly suited for occipital stimulation, reaching current densities of up to 0.089 A/m2. Compared to the Cz/Oz montage, our approach is also less likely to elicit phosphenes (Zaehle et al., 2010). It may be that our electrode montage (positions P7 and P8 according to the International 10-20 system), which differed from the montage used by Ruhnau et al. (2016) (Cz and Oz), was ineffective in modulating the SSR. One reason for this could be the propensity of the Cz/Oz montage to reach more medial occipital areas (Neuling et al. 2012) and hence the calcerine sulcus, from whence we reason the SSRs to originate. Therefore, anterior/superior—posterior/inferior tACS stimulation could affect neural firing in occipital cortex differently from left—right stimulation employed here. However, we believe the orientation of the calcarine sulcus alone cannot account for differential effects of different tACS directions. One reason is that SSRs do not have exclusively occipital sources. In fact, functional magnetic resonance imaging results demonstrate the involvement of a variety of widespread cortical regions, including frontal and prefrontal cortices (Ding et al., 2006; Di Russo et al., 2007; Pastor et al., 2003; Srinivasan et al., 2007, Li et al., 2015). It was also theorised that the SSR eliciting stimulus activates primary visual cortex directly through thalamocortical inputs, while more distant regions are recruited through indirect connections, although it remains possible that thalamic inputs lead signals directly to more anterior regions (Tononi et al., 1998). We therefore are not aware of any theoretical reason to expect one montage to work and not the other. Lastly, our choice of montage was not arbitrary, but based on modelling results by Neuling et al. (2012b).

### 4.3 Limitations

Interestingly, tACS at alpha frequency not only affects neural oscillations during stimulation, but has aftereffects as well. While the mechanism behind these effects is disputed (Vossen et al., 2015), it is possible that they led to carry over effects as we proceeded from one condition to the next. Another possible confound are skin sensations induced by tACS. More sensitive participants may better feel the stimulation and hence be able to distinguish between the experimental conditions. However, we controlled for this in three ways: Firstly, we gradually faded stimulation in and out, leading to sensation mostly confined to time periods outside the main trials. Secondly, the stimulation frequencies were so close together that participants were unable to discriminate between them, as indexed by subjective report (data not shown). Lastly, in the sham condition, we also faded in and out a weak tACS current at a frequency between the fast and slow, leading to the same skin sensations as are experienced during non-sham trials.

Yet another concession to make is the sample of *n* = 20. While this sample size is standard for tACS publications (average *n* = 17 in 50 studies reviewed in Table 1 by Schutter & Wischnewski, 2016), caution is indicated: Veniero et al. (2017) showed an effect of tACS on a perceptual estimation task in a sample of *n* = 19. However, trying to replicate the same effect in an independent sample in the very same study revealed only a null result at *n* = 20 as well as in the combined sample of *n* = 49. It is hence possible that there is actually a small effect, only that it is obscured by the spread of the data. However, we are reasonably confident in the validity of our null result, also because of our use of Bayesian statistics and because we could not even observe a trend. Still, given the large inter-subject variability in response to tACS (review in Krause & Cohen Kadosh, 2014), this is a possibility we cannot dismiss.

Another issue to consider is that the alpha frequency band (7.5 – 12.5 Hz), into which our SSR fell, may not easily be susceptible to tACS modulation. In particular, there may be a task and baseline power dependence, such that tACS can only modulate alpha when endogenous alpha is low: indeed, phase coherence between tACS at individual alpha frequency and endogenous brain oscillations over occipital pole have been reported to increase only when participants had their eyes open (Ruhnau et al., 2016b). Also, Neuling et al. (2013) tried to entrain alpha power while participants had their eyes either open or closed. Again, when eyes were closed and endogenous alpha power was high, tACS was unable to modulate it, possibly due to ceiling effects in the current densities: Neuling et al. (2012b) demonstrated that 417 μV/mm electric field can be induced with tACS over occipital regions; occipital alpha in awake monkeys has been estimated at 400 μV/mm (Bollimunta et al., 2008). When eyes are closed, alpha is expected to far exceed this value, hence there may not be enough tACS current strength available to modulate it. SSRs are known to be high in alpha power (Herrmann, 2001), especially in participants with already high endogenous alpha (Pigeau & Frame, 1992). However, the argument that SSRs may already be saturated is not convincing, given that the SSRs during suppressed periods in experiment 1 should have been substantially weaker than the dominant based on the past literature (Brown & Norcia 1997, Tononi et al 1998, Zhang et al 2010), implying that they were not yet at ceiling. Also in the CFS experiments, the target was small, hence it is even more unlikely to saturate SSR.

Another limitation concerns the specific choice of stimulus to investigate the causal role of the SSR in conscious perception: Both facilitation and adaptation of neural activity have been proposed as integral parts in models of binocular rivalry (Wilson, 2007) and CFS (Shimaoka & Kaneko, 2011). It is at least plausible that tACS itself causes both adaptation and facilitation of the neural activity. If so, then any effect on these paradigms might be due to these mechanisms, hence may be independent of a modulation of the SSR. If tACS affects both adaptation and facilitation symmetrically, then this would entail that our hypotheses could be formulated as two-tailed and that the observed null could be due to the opposing effects on adaptation and facilitation cancelling each other out. Alas, this concern cannot be ruled out by our control conditions.

Finally, we did not control for contrast in experiments 2 and 3. Hence, it is possible that had we titrated the stimulus contrast, we may have been able to demonstrate a small but consistent effects. In support of this, behavioural results in *figure* 6 suggests that detection was sometimes saturated in some subjects. In the future, one ought to control contrast through a QUEST procedure to ensure that all participants’ performance is 75% of 80% at the baseline.

## 5. Conclusion

Considering the above, one account for the absence of perceptual effects of tACS in the present study could be that the SSRs amplitude was too large to be significantly modulated by tACS. Without this modulation, there naturally would not be a behavioural effect. Again, future studies should concurrently record EEG to test if the SSR actually was modulated.

Since our results did not demonstrate a causal role of the SSR in conscious perception, the question of its causal role remains elusive. In the case of fMRI evidence pointing towards the role of the parietal cortex in bistability, such a causal role was supported by use of transcranial magnetic stimulation (Carmel et al., 2010; Kanai et al., 2010, 2011; Zarektskaya et al., 2010, 2013; Wood & Schauer et al., in prep). Also neural oscillations in the gamma band, which have been associated to binocular rivalry (Engel et al., 2001), have been modulated by tACS to also affect perception by disturbing inter-hemispheric phase coherence by use of two tACS montages, one on each hemisphere (Strüber et al., 2013). This leads to another issue that must be addressed: if endogenous alpha has already some causal role, then modulating an alpha based SSR through tACS will by necessity also modulate endogenous alpha. Hence, with any effect, we cannot be sure if what we observe is due to a modulation of endogenous alpha or the SSR, since brain oscillations are not independent of the signal designed to probe them (Keitel et al., 2014). In the worst case, both might have opposing effects that cancel each other out.

In conclusion, it is still doubtful if SSRs have a causal role in conscious perception. Future studies should record EEG data concurrently with tACS in order to test whether SSRs are actually modulated while perceptual measures are taken. Given the large inter-subject variability observed with tACS, this would also allow investigating if EEG parameters predict the direction and strength of the tACS effect. For now, we tentatively conclude that the SSR merely correlates with the content of visual perception, however, its modulation does not affect perception.

## Acknowledgements

GS and AB conceptualised and designed the study. NT and GS coded the visual stimuli. CO collected and preprocessed the data. GS performed the data analysis. CO and GS drafted the manuscript, which received critical revisions from NT and AB. All authors approved of the final version of the manuscript for submission. We thank Julian Rodney Matthews for his aid in writing the preprocessing scripts. This work was supported by the Barbara-Wengeler Foundation.

## References

Andersen, S. K., Müller, M. M., & Hillyard, S. A. (2011). Tracking the allocation of attention in visual scenes with steady-state evoked potentials. Cognitive neuroscience of attention, 2, 197–216.

Andersen, S. K., Müller, M. M., & Hillyard, S. A. (2015). Attentional selection of feature conjunctions is accomplished by parallel and independent selection of single features. Journal of Neuroscience, 35(27), 9912–9919.

Antal, A., & Paulus, W. (2013). Transcranial alternating current stimulation (tACS). Frontiers in human neuroscience, 7.

Battleday, R. M., Muller, T., Clayton, M. S., & Kadosh, R. C. (2014). Mapping the mechanisms of transcranial alternating current stimulation: a pathway from network effects to cognition. Frontiers in psychiatry, 5.

Blake, R., & Logothetis, N. (2002). Visual competition. Nature Reviews Neuroscience, 3, 13–21.

Bollimunta, A., Chen, Y., Schroeder, C. E., & Ding, M. (2008). Neuronal mechanisms of cortical alpha oscillations in awake-behaving macaques. Journal of Neuroscience, 28(40), 9976–9988.

Bor, D., Schwartzman, D. J., Barrett, A. B., & Seth, A. K. (2017). Theta-burst transcranial magnetic stimulation to the prefrontal or parietal cortex does not impair metacognitive visual awareness. PloS one, 12(2), e0171793.

Brown, R. J., & Norcia, A. M. (1997). A method for investigating binocular rivalry in real-time with the steady-state VEP. Vision research, 37(17), 2401–2408.

Busch, N. A., Dubois, J., & VanRullen, R. (2009). The phase of ongoing EEG oscillations predicts visual perception. Journal of Neuroscience, 29(24), 7869–7876.

Carmel, D., Walsh, V., Lavie, N., & Rees, G. (2010). Right parietal TMS shortens dominance durations in binocular rivalry. Current biology, 20(18), R799–R800.

Di Russo, F., Pitzalis, S., Aprile, T., Spitoni, G., Patria, F., Stella, A., & Hillyard, S. A. (2007). Spatiotemporal analysis of the cortical sources of the steady-state visual evoked potential. Human brain mapping, 28(4), 323–334.

Ding, J., Sperling, G., & Srinivasan, R. (2005). Attentional modulation of SSVEP power depends on the network tagged by the flicker frequency. Cerebral cortex, 16(7), 1016–1029.

Engel, A. K., Fries, P., & Singer, W. (2001). Dynamic predictions: oscillations and synchrony in top–down processing. Nature Reviews Neuroscience, 2(10), 704–716.

Fleming, S. M., & Lau, H. C. (2014). How to measure metacognition. Frontiers in human neuroscience, 8.

Herrmann, C. S. (2001). Human EEG responses to 1–100 Hz flicker: resonance phenomena in visual cortex and their potential correlation to cognitive phenomena. Experimental brain research, 137(3-4), 346–353.

Herrmann, C. S., Rach, S., Neuling, T., & Strüber, D. (2013). Transcranial alternating current stimulation: a review of the underlying mechanisms and modulation of cognitive processes. Frontiers in human neuroscience, 7, 279.

Jiang, Y., Costello, P., & He, S. (2006). Processing of invisible stimuli: faster for upright faces and recognizable words to overcome interocular suppression. Psychol. Sci.

Kanai, R., Bahrami, B., & Rees, G. (2010). Human parietal cortex structure predicts individual differences in perceptual rivalry. Current Biology, 20(18), 1626–1630.

Kanai, R., Carmel, D., Bahrami, B., & Rees, G. (2011). Structural and functional fractionation of right superior parietal cortex in bistable perception. Current biology, 21(3), R106–R107.

Kanai, R., Chaieb, L., Antal, A., Walsh, V., & Paulus, W. (2008). Frequency-dependent electrical stimulation of the visual cortex. Current Biology, 18(23), 1839–1843.

Keitel, C., Quigley, C., & Ruhnau, P. (2014). Stimulus-driven brain oscillations in the alpha range: entrainment of intrinsic rhythms or frequency-following response?. Journal of Neuroscience, 34(31), 10137–10140.

Krause, B., & Kadosh, R. C. (2014). Not all brains are created equal: the relevance of individual differences in responsiveness to transcranial electrical stimulation. Frontiers in systems neuroscience, 8(25), 1–12.

Laczó, B., Antal, A., Niebergall, R., Treue, S., & Paulus, W. (2012). Transcranial alternating stimulation in a high gamma frequency range applied over V1 improves contrast perception but does not modulate spatial attention. Brain stimulation, 5(4), 484–491.

Lawwill, T., & Biersdorf, W. R. (1968). Binocular rivalry and visual evoked responses. Investigative Ophthalmology & Visual Science, 7(4), 378–385.

Levelt, W. J. (1967). Note on the distribution of dominance times in binocular rivalry. British Journal of Psychology, 58(1-2), 143–145.

Li, F., Tian, Y., Zhang, Y., Qiu, K., Tian, C., Jing, W., & Xu, P. (2015). The enhanced information flow from visual cortex to frontal area facilitates SSVEP response: evidence from model-driven and data-driven causality analysis. Scientific reports, 5.

Matthews, J., Schröder, P., Kaunitz, L., van Boxtel, J. J., & Tsuchiya, N. (2018). Conscious access in the near absence of attention: critical extensions on the dual-task paradigm. Philosophical Transactions of the Royal Society B: Biological Sciences, 373(1755), 20170352.

Matthews, J., Wu, J., Corneille, V., Hohwy, J., van Boxtel, J., & Tsuchiya, N. (2019). Sustained conscious access to incidental memories in RSVP. Attention, Perception, & Psychophysics, 81(1), 188–204.

Moliadze, V., Atalay, D., Antal, A., & Paulus, W. (2012). Close to threshold transcranial electrical stimulation preferentially activates inhibitory networks before switching to excitation with higher intensities. Brain stimulation, 5(4), 505–511.

Montemurro, M. A., Rasch, M. J., Murayama, Y., Logothetis, N. K., & Panzeri, S. (2008). Phase-of-firing coding of natural visual stimuli in primary visual cortex. Current Biology, 18(5), 375–380.

Neuling, T., Rach, S., & Herrmann, C. S. (2013). Orchestrating neuronal networks: sustained after-effects of transcranial alternating current stimulation depend upon brain states. Frontiers in human neuroscience, 7.

Neuling, T., Rach, S., Wagner, S., Wolters, C. H., & Herrmann, C. S. (2012a). Good vibrations: oscillatory phase shapes perception. Neuroimage, 63(2), 771–778.

Neuling, T., Wagner, S., Wolters, C. H., Zaehle, T., & Herrmann, C. S. (2012b). Finite-element model predicts current density distribution for clinical applications of tDCS and tACS. Frontiers in psychiatry, 3.

Norcia, A. M., Appelbaum, L. G., Ales, J. M., Cottereau, B. R., & Rossion, B. (2015). The steady-state visual evoked potential in vision research: a review. Journal of vision, 15(6), 4–4.

Pastor, M. A., Artieda, J., Arbizu, J., Valencia, M., & Masdeu, J. C. (2003). Human cerebral activation during steady-state visual-evoked responses. Journal of neuroscience, 23(37), 11621–11627.

Persaud, N., McLeod, P., & Cowey, A. (2007). Post-decision wagering objectively measures awareness. Nature neuroscience, 10(2), 257–261.

Pigeau, R. A., & Frame, A. M. (1992). Steady-state visual evoked responses in high and low alpha subjects. Electroencephalography and Clinical Neurophysiology/Evoked Potentials Section, 84(2), 101–109.

Pogosyan A., Gaynor L. D., Eusebio A., Brown P. (2009) Boosting cortical activity at Beta-band frequencies slows movement in humans. Curr Biol 19: 1637–1641.

Polanía, R., Nitsche, M. A., Korman, C., Batsikadze, G., & Paulus, W. (2012). The importance of timing in segregated theta phase-coupling for cognitive performance. Current Biology, 22(14), 1314–1318.

Reato, D., Rahman, A., Bikson, M., & Parra, L. C. (2010). Low-intensity electrical stimulation affects network dynamics by modulating population rate and spike timing. Journal of Neuroscience, 30(45), 15067–15079.

Regan D. (1989). Human brain electrophysiology: evoked potentials and evoked magnetic fields in science and medicine. New York: Elsevier.

Regan, D. (1966). An effect of stimulus colour on average steady-state potentials evoked in man. Nature, 210, 1056–1057.

Rouder, J. N., Morey, R. D., Speckman, P. L., & Province, J. M. (2012). Default Bayes factors for ANOVA designs. Journal of Mathematical Psychology, 56(5), 356–374.

Rounis, E., Maniscalco, B., Rothwell, J. C., Passingham, R. E., & Lau, H. (2010). Theta-burst transcranial magnetic stimulation to the prefrontal cortex impairs metacognitive visual awareness. Cognitive neuroscience, 1(3), 165–175.

Ruhnau, P., Keitel, C., Lithari, C., Weisz, N., & Neuling, T. (2016a). Flicker-driven responses in visual cortex change during matched-frequency transcranial alternating current stimulation. Frontiers in human neuroscience, 10.

Ruhnau, P., Neuling, T., Fuscá, M., Herrmann, C. S., Demarchi, G., & Weisz, N. (2016b). Eyes wide shut: transcranial alternating current stimulation drives alpha rhythm in a state dependent manner. Scientific reports, 6, 27138.

Schutter, D. J., & Wischnewski, M. (2016). A meta-analytic study of exogenous oscillatory electric potentials in neuroenhancement. Neuropsychologia, 86, 110–118.

Sejnowski, T. J., & Paulsen, O. (2006). Network oscillations: emerging computational principles. Journal of Neuroscience, 26(6), 1673–1676.

Sherman, M. T., Barrett, A. B., & Kanai, R. (2015). Inferences about consciousness using subjective reports of confidence. In: Overgaard, Morten (ed.) Behavioral Methods in Consciousness Research. Oxford University Press, Oxford, UK.

Sherman, M. T., Kanai, R., Seth, A. K., & VanRullen, R. (2016). Rhythmic influence of top–down perceptual priors in the phase of prestimulus occipital alpha oscillations. Journal of cognitive neuroscience 28(9), 1318–1330.

Shimaoka, D., & Kaneko, K. (2011). Dynamical systems modeling of continuous flash suppression. Vision Research, 51(6), 521–528.

Spaak, E., de Lange, F. P., & Jensen, O. (2014). Local entrainment of alpha oscillations by visual stimuli causes cyclic modulation of perception. Journal of Neuroscience, 34(10), 3536–3544.

Srinivasan, R., Fornari, E., Knyazeva, M. G., Meuli, R., & Maeder, P. (2007). fMRI responses in medial frontal cortex that depend on the temporal frequency of visual input. Experimental brain research, 180(4), 677–691.

Strüber, D., Rach, S., Trautmann-Lengsfeld, S. A., Engel, A. K., & Herrmann, C. S. (2014). Antiphasic 40 Hz oscillatory current stimulation affects bistable motion perception. Brain topography, 27(1), 158–171.

Thut, G., Schyns, P. G., & Gross, J. (2011). Entrainment of perceptually relevant brain oscillations by non-invasive rhythmic stimulation of the human brain. Frontiers in psychology, 2.

Tononi, G., Srinivasan, R., Russell, D. P., & Edelman, G. M. (1998). Investigating neural correlates of conscious perception by frequency-tagged neuromagnetic responses. Proceedings of the National Academy of Sciences, 95(6), 3198–3203.

Tsuchiya, N., & Koch, C. (2005). Continuous flash suppression reduces negative afterimages. Nature neuroscience, 8(8), 1096–1101.

VanRullen, R., Busch, N. A., Drewes, J., & Dubois, J. (2011). Ongoing EEG phase as a trial-by-trial predictor of perceptual and attentional variability. Frontiers in psychology, 2(60), 1–9.

Vialatte, F. B., Maurice, M., Dauwels, J., & Cichocki, A. (2010). Steady-state visually evoked potentials: focus on essential paradigms and future perspectives. Progress in neurobiology, 90(4), 418–438.

Vossen, A., Gross, J., & Thut, G. (2015). Alpha power increase after transcranial alternating current stimulation at alpha frequency (α-tACS) reflects plastic changes rather than entrainment. Brain stimulation, 8(3), 499–508.

Wilson, H. R. (2007). Minimal physiological conditions for binocular rivalry and rivalry memory. Vision research, 47(21), 2741–2750.

Witkowski, M., Garcia-Cossio, E., Chander, B. S., Braun, C., Birbaumer, N., Robinson, S. E., & Soekadar, S. R. (2016). Mapping entrained brain oscillations during transcranial alternating current stimulation (tACS). Neuroimage, 140, 89–98.

Yang, E., Brascamp, J., Kang, M. S., & Blake, R. (2014). On the use of continuous flash suppression for the study of visual processing outside of awareness. Frontiers in psychology, 5.

Zaehle, T., Rach, S., & Herrmann, C. S. (2010). Transcranial alternating current stimulation enhances individual alpha activity in human EEG. PloS one, 5(11), e13766.

Zaretskaya, N., Anstis, S., & Bartels, A. (2013). Parietal cortex mediates conscious perception of illusory gestalt. Journal of Neuroscience, 33(2), 523–531.

Zaretskaya, N., Thielscher, A., Logothetis, N. K., & Bartels, A. (2010). Disrupting parietal function prolongs dominance durations in binocular rivalry. Current biology, 20(23), 2106–2111.

Zhang, P., Jamison, K., Engel, S., He, B., & He, S. (2011). Binocular rivalry requires visual attention. Neuron, 71(2), 362–369.

Zizlsperger, L., Kümmel, F., & Haarmeier, T. (2016). Metacognitive confidence increases with, but does not determine, visual perceptual learning. PloS one, 11(3), e0151218.

